## Supplementary material for "*Trans*-eQTL hotspots shape complex traits by modulating cellular states"

#### Supplementary material - Table of contents

| *Section name* | *Pages* |
|---|---|
| 1. Supplementary text | 1- 4 |
| 2. Supplementary figures | 5- 17 |
| 3. Supplementary tables | 18- 19 |
| 4. Supplementary references | 20- 21 |

### Supplementary text

*Genetic correlations are not due to factors that are shared across traits*

The growth traits analyzed here are not entirely independent from each other, as reflected by significant correlations among them (Supplementary Figure 1; see also Bloom et al. [1]). This raises the possibility that the genetic correlations observed for the 46 traits may reflect the same underlying biological relationships, rather than trait-specific connections. To test this, we explored whether the genetic correlations for different traits were driven by factors that are shared among the 46 traits.

Growth traits were measured as colony sizes on solid plates with either YNB or YPD agar medium (Table S1). This means that genetic influences on how well a segregant grows on these two media could also influence all traits measured on the respective medium. Such shared growth effects could be reflected in expression-trait correlations that recur across traits. To test for the influence of the solid agar medium, we regressed each trait on growth on YNB or YPD (Table S1) and calculated correlations between gene expression and the residual traits. As expected, this treatment eliminated all correlations for YNB and YPD (Supplementary Figure 2). For three traits, more than half of the correlations lost significance (galactose: 90%, 4-Hydroxybenzaldehyde: 89%, 4-Nitroquinoline oxide: 68%). For the remaining 41 traits, a median of only 11% of correlations became non-significant (Supplementary Figure 2). Thus, most correlations between gene expression and growth do not simply reflect growth on the given solid medium but instead reflect trait-specific biology.

To explore whether unknown shared factors beyond the solid medium could account for covariation across conditions, we performed a principal component analysis on the growth traits. Any strong shared factors shaping growth across conditions would be reflected in large amounts of variation attributed to the first few principal components. Instead, the first principal component accounted for just 16.5% of variance among 42 traits (these analyses excluded 4 traits with high missing data; Table S1). Cumulatively, the first five principal components explained only 42% of variance (Supplementary Figure 3). Together, these analyses show that there are no strong common factors that drive most of the variation among traits, and that most of the correlations between gene expression and growth traits are specific to each trait.

*Influence of sample size on ability to detect genetic correlations*

Most of the genetic correlations between gene expression and traits had fairly modest magnitudes, with an overall median absolute correlation coefficient of 0.11. We hypothesized that our ability to detect thousands of correlations with such modest magnitudes with high statistical significance was possible due to the large size of the segregant panel. To test the influence of sample size, we subsampled the 979 segregants to smaller panels of 250, 500 and 750 segregants and recomputed the correlations. As expected, the number of significant (5% FDR) correlations increased with larger segregant panels (Supplementary Figure 5). At the same time, the median magnitude of the detected significant associations decreased with increasing sample size (Supplementary Figure 6), as expected for larger samples that can detect correlations of weaker magnitude. Hence, the size of the segregant panel permits discovery of thousands of associations between gene expression and genetically complex traits.

*Causal models underlying overlapping vs colocalized local eQTLs and gQTLs*

All 591 gQTLs in our dataset overlapped with at least one local eQTL. This overlap can arise from several scenarios (Figure 2A). First, under mediation or "vertical pleiotropy" [2,3], causal DNA variants alter gene expression in the baseline condition, which then affects growth when segregants encounter the given environmental condition. Second, under "horizontal pleiotropy", causal DNA variants also affect gene expression and growth but do so independently, through distinct pathways. Third, overlapping QTLs can arise from distinct, linked causal variants in physical proximity that affect only gene expression or only growth. QTLs with shared, pleiotropic variants are called "colocalized" to distinguish them from QTLs that overlap due to simple proximity between distinct causal variants [4]. To distinguish between these scenarios, we performed colocalization tests [5], which ask if a model of shared, pleiotropic variants can be rejected in favor of a model with different causal variants for gene expression and growth (Figure 2A). We tested 2,074 pairs of strong (both logarithm of the odds [LOD] scores ≥ 10) local eQTLs and gQTLs. These pairs comprised 188 gQTLs from 45 conditions that overlapped with at least one local eQTL for one of 581 genes. The pleiotropic model was not rejected at about half (1,052) of the QTL pairs. At these QTL pairs, which included 95% (178 / 188) of the analyzed gQTLs (Figure 2B & C), the same DNA variants may cause both the local eQTL and the gQTL.

*Local eQTLs at gQTLs with known causal genes*

We observed that 87% of the gQTLs were colocalized with local eQTLs for multiple genes. To gain intuition about potential causality of these colocalized local eQTLs, we examined gQTLs at which a gene has been demonstrated to be causal experimentally or is highly likely to be causal based on gene function.

We examined a gQTL for growth in presence of lithium chloride that contains the gene *ENA1*, which encodes a membrane-bound pump that controls efflux of lithium ions from the cell (Supplementary Figure 7A). In yeast, variable copy number at the *ENA* locus underlies growth variation linked to this locus [6,7]. Three local eQTLs overlapped with this gQTL, two of which (affecting *ENA1* and *HEM13*) were classified as colocalized. The *ENA1* local eQTL is extremely strong (LOD = 359, r = -0.89), as might be expected if higher *ENA1* expression from the BY allele is caused by a higher number of expressed copies in the BY compared to the RM strain. While we do not know the structure and gene copy numbers in the ENA locus in the parental BY and RM strains of this cross, it is reasonable to assume that *ENA1* is the causal gene at this gQTL. This example suggests that colocalization analyses can detect cases of causal colocalization, especially when the underlying effects are strong.

A gQTL on chromosome VII that shapes growth in the presence of manganese sulfate is largely, and perhaps exclusively, caused by a single missense variant in the gene *PMR1* [8]. *PMR1* does not have a local eQTL. In our colocalization analyses, six of the eight local eQTLs for genes other than *PMR1* that overlap this gQTL were classified as colocalized with the gQTL (Supplementary Figure 7B). While we cannot rule out that the causal variants creating these six local eQTLs contribute minor effects to this gQTL in addition to the missense variant in *PMR1*, it seems likely that most of these eQTLs were incorrectly flagged as colocalized due to linkage with the causal missense variant in *PMR1*.

*Genetic correlations at gene / trait pairs with overlapping local eQTLs and gQTLs*

To ask whether local eQTLs may contribute to genetic correlations, we compared the magnitudes of genetic correlations at gene / trait pairs with and without overlapping local eQTLs and gQTLs. Genetic correlations were stronger for gene / trait pairs with local eQTL / gQTL overlap than for pairs with local eQTLs that did not overlap a gQTL (average absolute r = 0.095 vs 0.066; t-test $p < 2.2e-16$). Because eQTL / gQTL overlap can arise from shared pleiotropic variants or from distinct causal variants (Figure 2A), we divided gene / trait pairs with  eQTL / gQTL overlap into pairs classified as colocalized versus pairs at which colocalization was rejected. There was no difference in the strength of genetic correlations between these groups (p = 0.95), and there was no association between colocalization status and the presence of a significant genetic correlation (Fisher's exact test (FET): p = 0.3). Thus, local eQTLs contribute to genetic correlations when they overlap a detected gQTL. This signal is expected, given that overlapping QTLs arise from linked variants, creating a correlation between the affected traits. However this signal may be due to either shared causal variants or distinct but linked causal variants.

*Interpretation of GO enrichments among genes with genetic correlations*

GO enrichment results of genes with genetic correlations revealed some results that are not immediately intuitive, including correlations between higher growth in some of the 46 conditions and a gene expression signature associated with *slower* growth in chemostats (Figure 6). Interpreting exactly how a genetic predisposition for a given gene expression signature may impact growth in a given condition is challenging with the current data. The doses of the stressor used for phenotyping were sufficiently low to be sub-lethal for most segregants, permitting study of quantitative growth variation rather than binary survival. The end-point colony sizes studied here likely integrate rich but unmeasured growth dynamics (similar to that demonstrated in Li et al. [9], including how quickly cells began growing, their rates and duration of growth, and relative rates of cell division and cell death. For example, cells genetically predisposed to fast growth may be more likely to die in a certain condition, such that the resulting colony is mainly formed by slowly growing cells. Larger colonies could also reflect more cells or larger cells. Finally, the environmental conditions were present throughout the incubation period, which may involve different cellular responses than an acute stress applied to the cell [10] or adaptive responses induced by the cell's acute stress response pathways in order to survive a prolonged exposure to the same stress [10,11].

### Supplementary figures

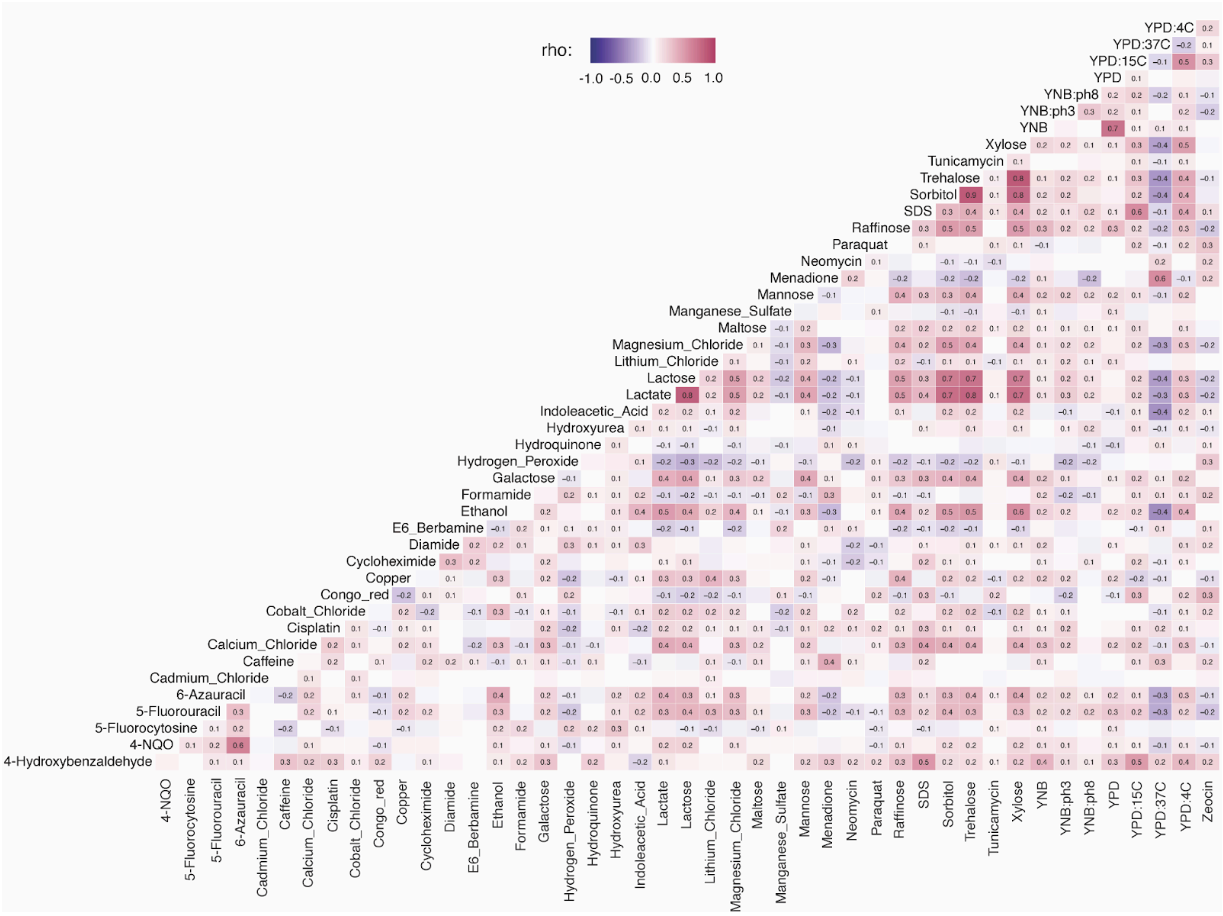


Supplementary Figure 1: Spearman correlation coefficients (rho) for all pairs of traits. Significant spearman correlation coefficients are displayed within the cells of the heatmap.

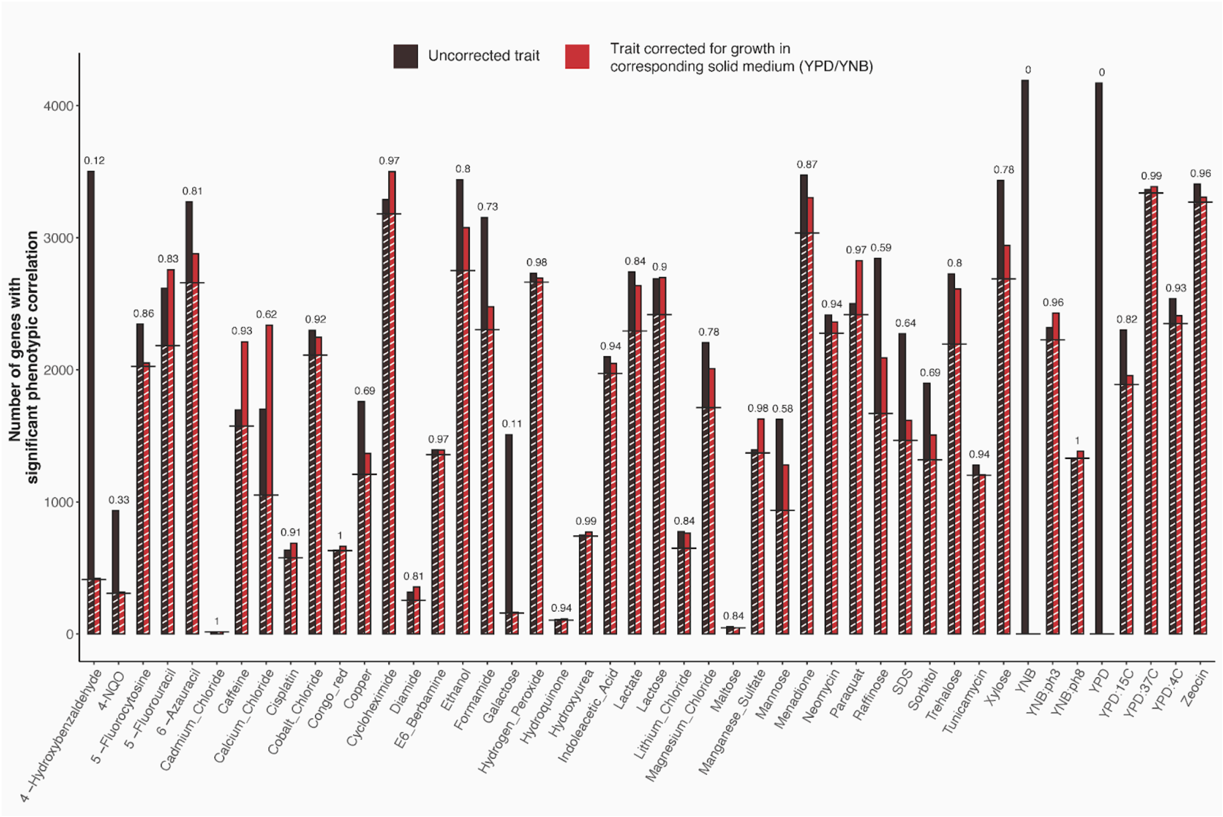


Supplementary Figure 2: Number of genes with significant genetic correlations at 5% FDR before (black) and after (red) correcting for growth on the respective solid medium (YPD or YNB). White shading and black horizontal lines show genes that have significant correlations before as well as after correction. Numbers above each pair of black and red bars indicate the fraction of genes with a genetic correlation that persisted after correction for the solid medium.

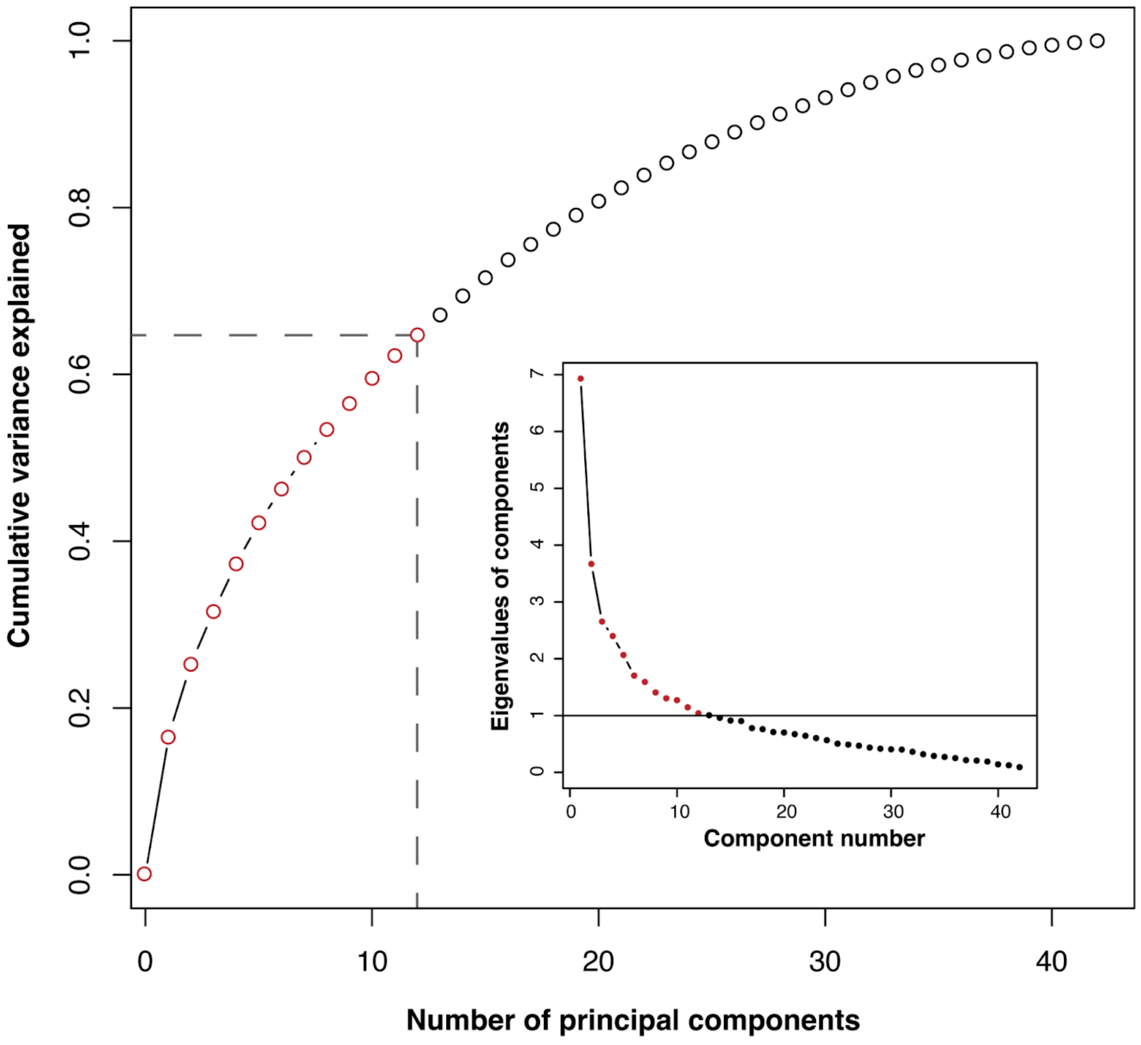


Supplementary Figure 3: Cumulative distribution of the proportion of variance among growth traits explained by principal components. The inset shows a scree plot for the same principal component analysis. The first 12 principal components account for most of the variance in growth traits based on the Kaiser criterion (Eigenvalue of component ≥ 1, points indicated in red) [12]. Together, these 12 components account for ~65% of the variance in growth traits.

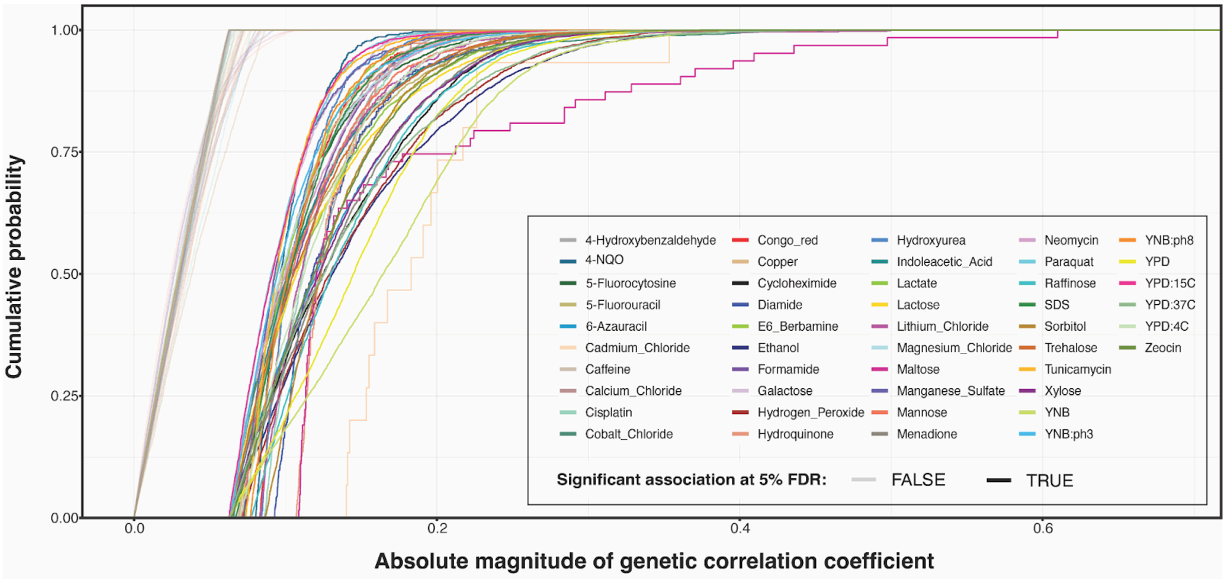


Supplementary Figure 4: Cumulative distribution of absolute magnitudes of correlation coefficients for genetic correlations between gene expression and growth in each of the 46 conditions. For each condition, the figure shows separate distributions for significant (FDR of 5%) correlations (curves in saturated colors) and non-significant correlations (curves in pale colors).

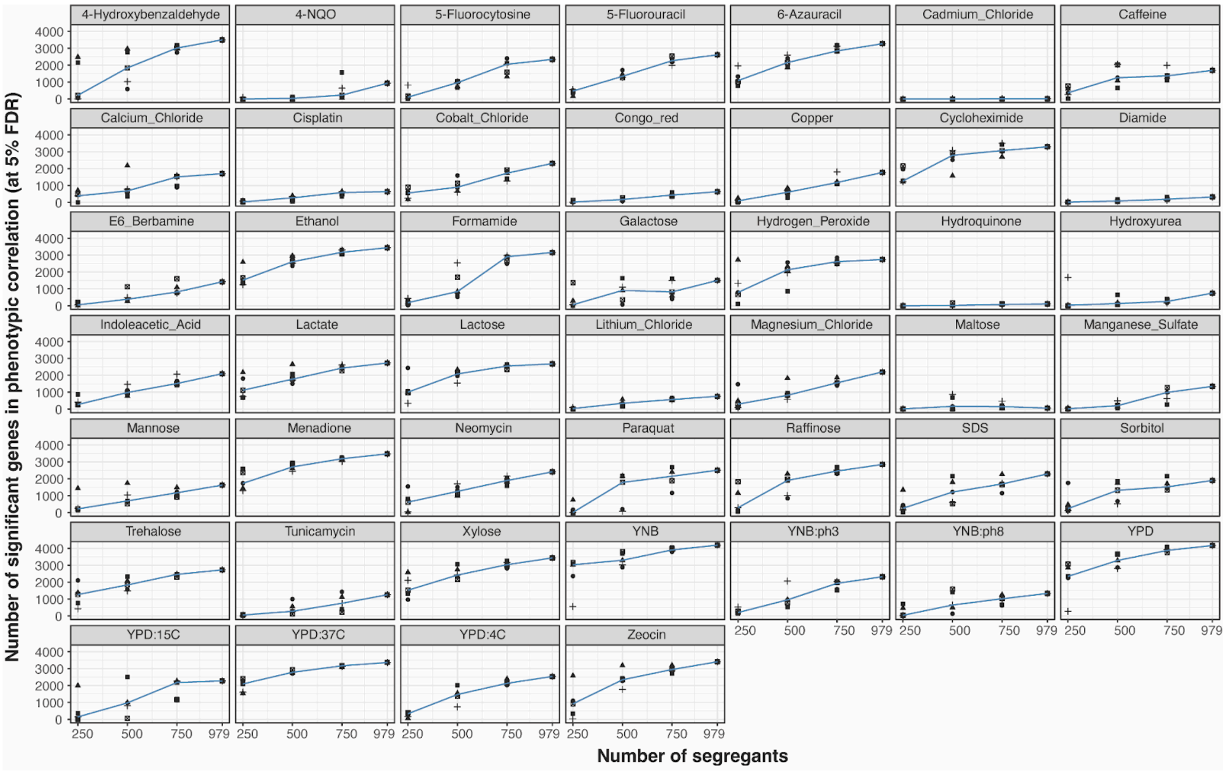


Supplementary Figure 5: The number of genes with significant correlation between expression and growth in different conditions as a function of sample size. We performed five random draws per sample size, indicated by different symbols. The trend line represents the median number of genes with significant correlation across the five draws for a given sample size.

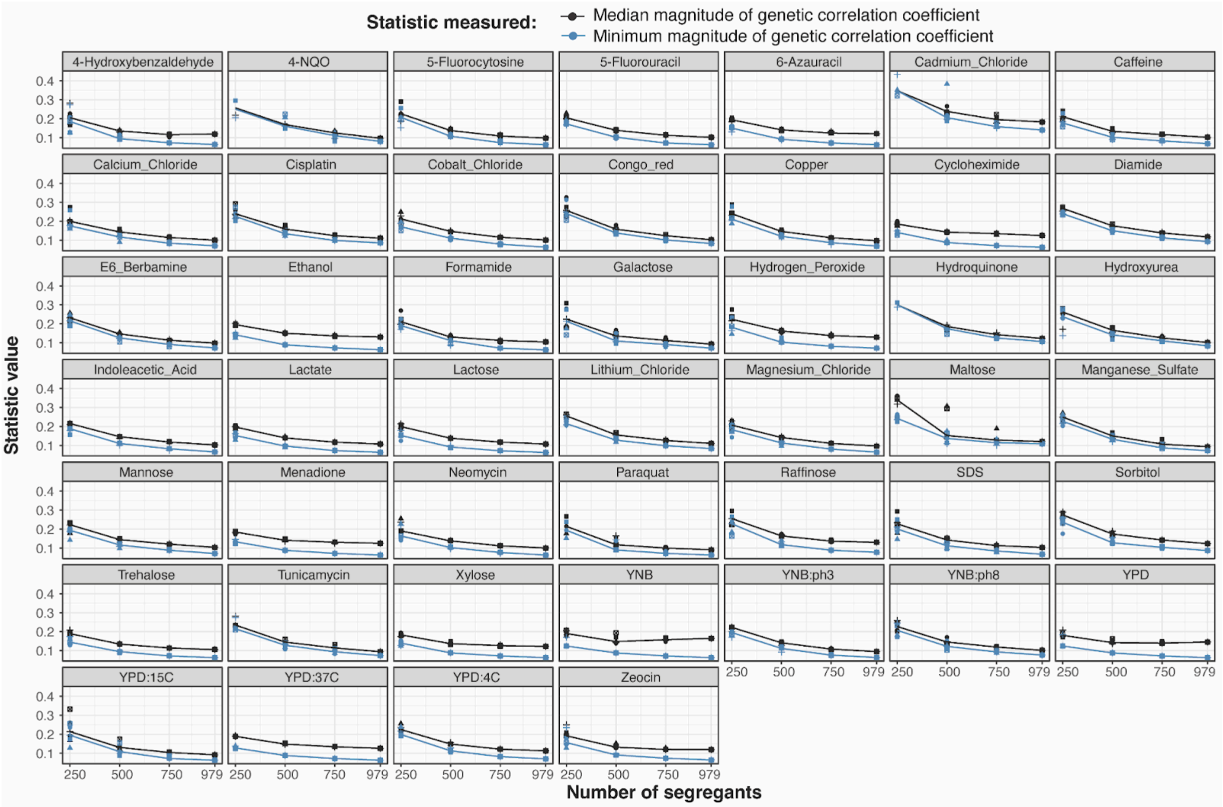


Supplementary Figure 6: Medians and minimums of the magnitudes of the correlation coefficients for significant correlations between gene expression and growth as a function of sample size. We performed five random draws per sample size, indicated by different symbols. The trend line represents the median value of the statistic across the five draws for a given sample size.

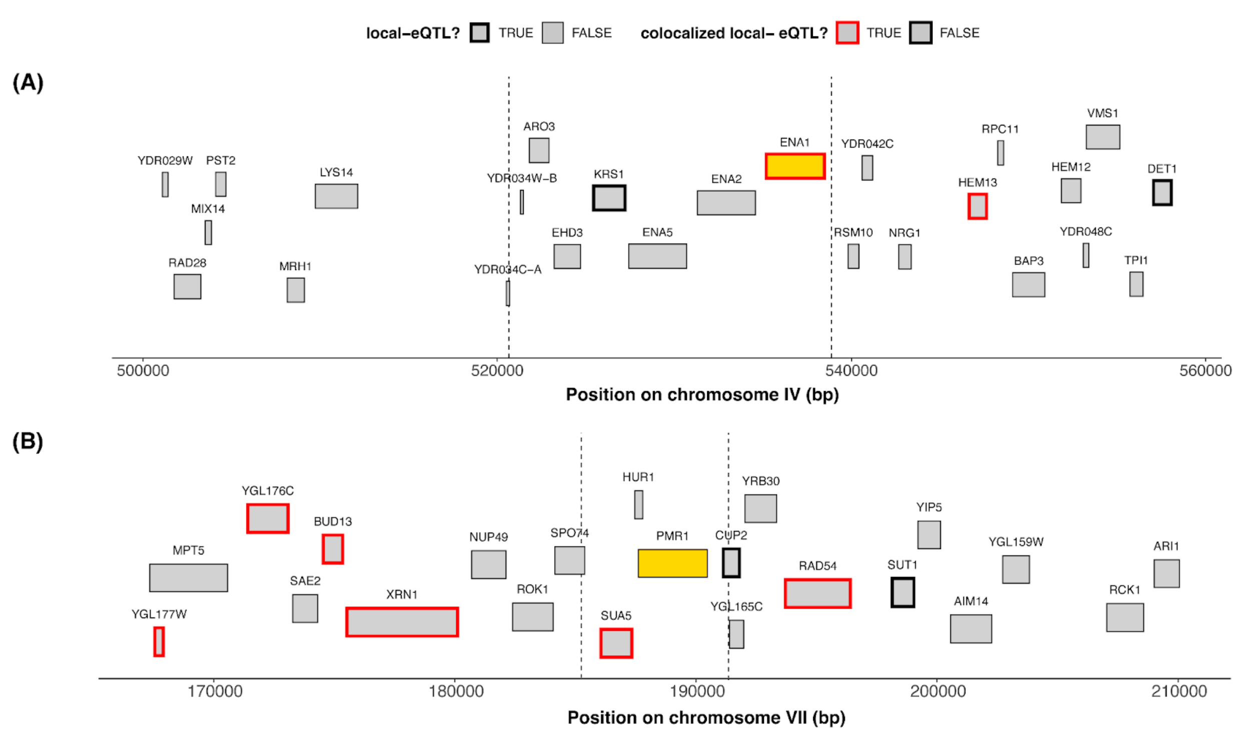


Supplementary Figure 7: Examples of local eQTLs at gQTLs for which the causal gene was either demonstrated experimentally (*PMR1*) or is very likely based on gene function (the *ENA* locus). The two panels show chromosome regions. Genes are shown as boxes. Causal genes for the given gQTL are shown as yellow boxes. Dotted vertical lines represent the 95% confidence intervals of gQTL location. Genes that have a local eQTL with LOD ≥ 10 and with a confidence interval that overlaps the gQTL are indicated by bold outlines, while genes that do not have a local eQTL are shown with thin outlines. Genes whose local eQTL is colocalized with the gQTL (the test for two separate QTLs was not significant; $p > 0.05$) are indicated by red outlines. Genes whose local eQTL is not colocalized with the gQTL (the test for two separate QTLs was significant at $p < 0.05$) are indicated by black outlines. (A) A gQTL for growth in the presence of lithium chloride. *ENA1* is the likely causal gene for this gQTL [6,7]. Note that *ENA1* is correctly flagged as having a colocalized eQTL, but so is the additional gene *HEM13*. (B) A gQTL for growth in the presence of manganese sulfate. A missense variant in *PMR1* has been experimentally shown to be causal for this gQTL [8]. Note that *PMR1* does not have a local eQTL and therefore cannot be detected by this colocalization analysis. Instead, the analysis flagged six other genes as having local eQTLs that are colocalized with this gQTL (red boxes). These genes are likely false positives due to linkage.

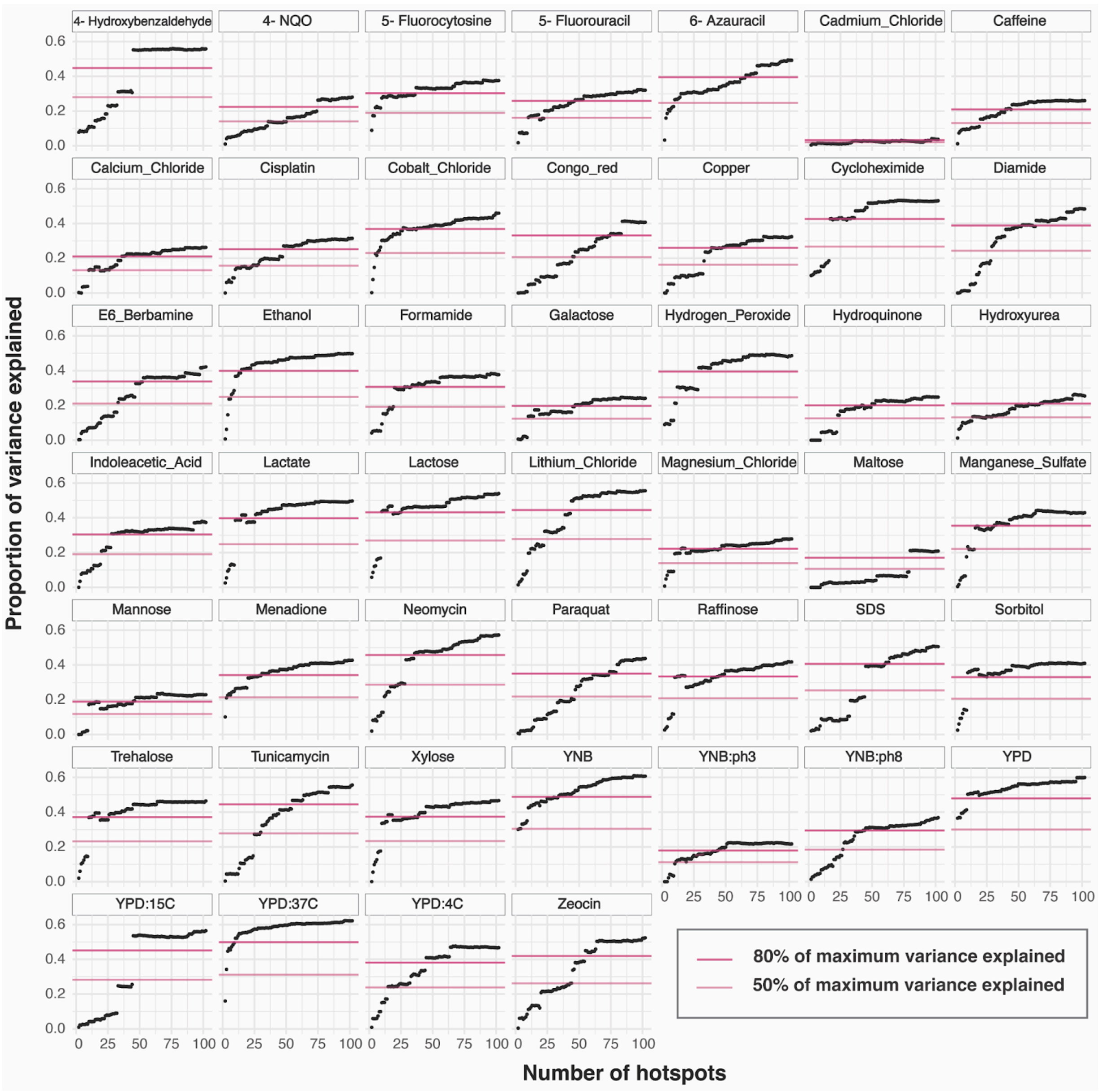


Supplementary Figure 8: Proportion of growth variance explained by top trans-eQTL hotspots, which are ranked from 2 to 102 according to the number of genes whose expression they affect, for each of the 46 growth conditions. The values corresponding to 80% and 50% of the maximum variance explained by the trans eQTL hotspot sets for each of the growth conditions are indicated by the dark pink and light pink lines, respectively.

146

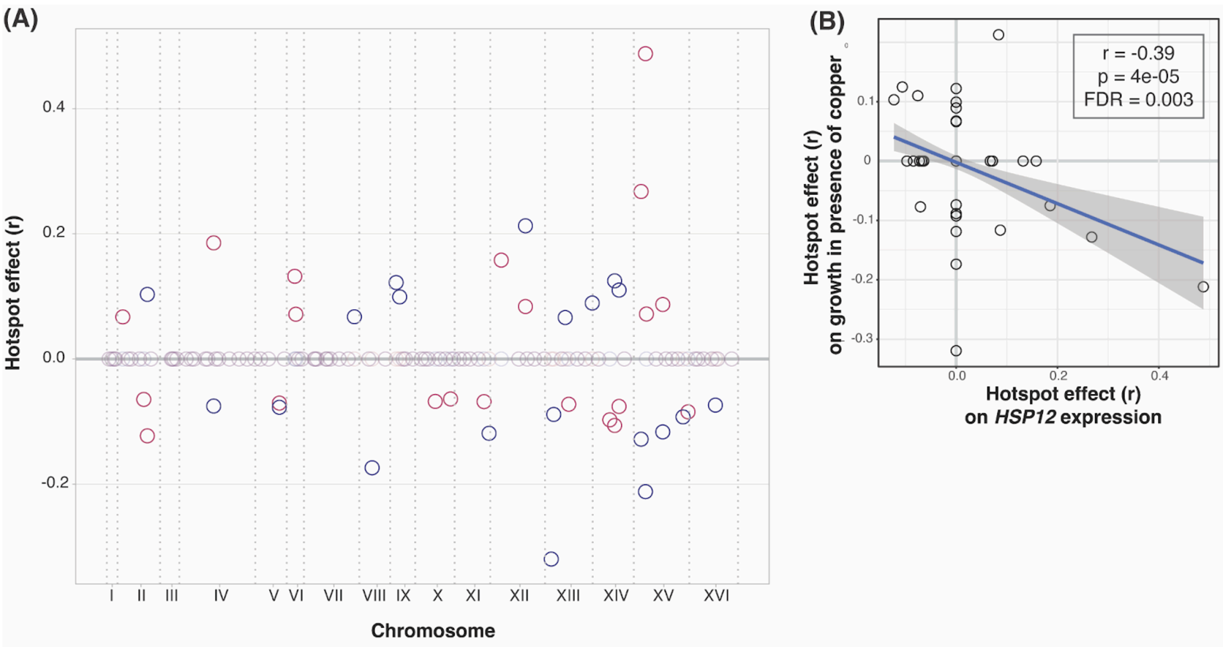


Supplementary Figure 9: Correlation of *trans*-eQTL hotspot effects on expression of *HSP12* and on growth in the presence of copper. (A) The plot shows the genomic locations of the *trans*-eQTL hotspots and their effect on the expression of *HSP12* (red circles) and on growth in the presence of copper (blue circles). Effects are coefficients of correlation between trait and genotype at the hotspot marker. Pale circles at the zero line indicate hotspot effects estimated as zero (Methods). (B) The scatterplot shows the effects of *trans*-eQTL hotspots on *HSP12* expression and on growth in the presence of copper, as shown in A. The regression line along with its 95% confidence interval is also displayed.

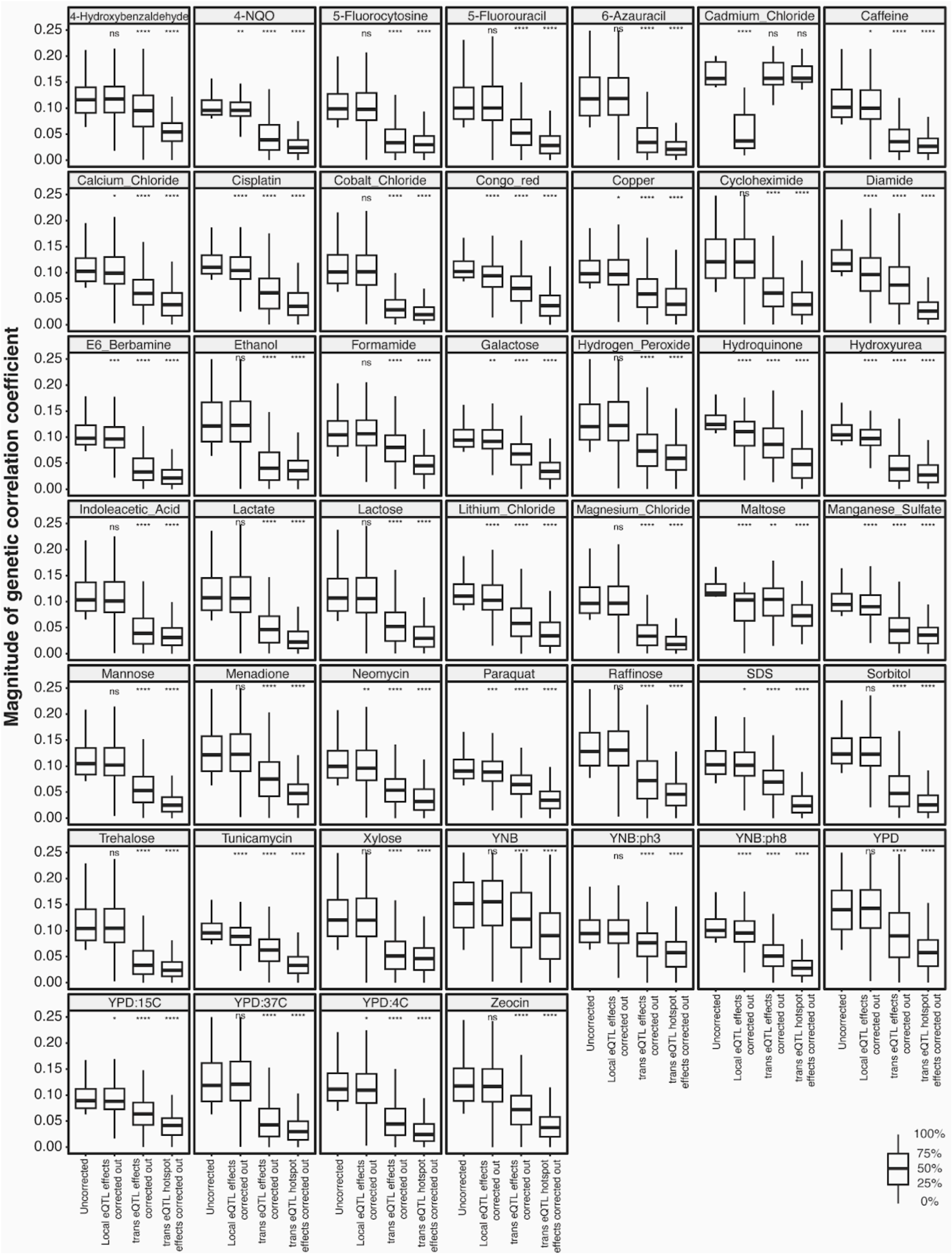


Supplementary Figure 10: Distributions of magnitudes of genetic correlation coefficients before and after correcting for the effects of local eQTLs, *trans*-eQTLs and *trans*-eQTL hotspots. Descriptions of the different categories are as in Figure 5C. The significance of the difference between the medians of the distributions with respect to the ‘uncorrected’ category was computed using the Wilcoxon test and significance indicated as follows: *ns* - not significant; * $0.01 < p < 0.05$; ** $p < 0.01$; *** $p < 0.001$; **** $p < 1e\text{-}04$.

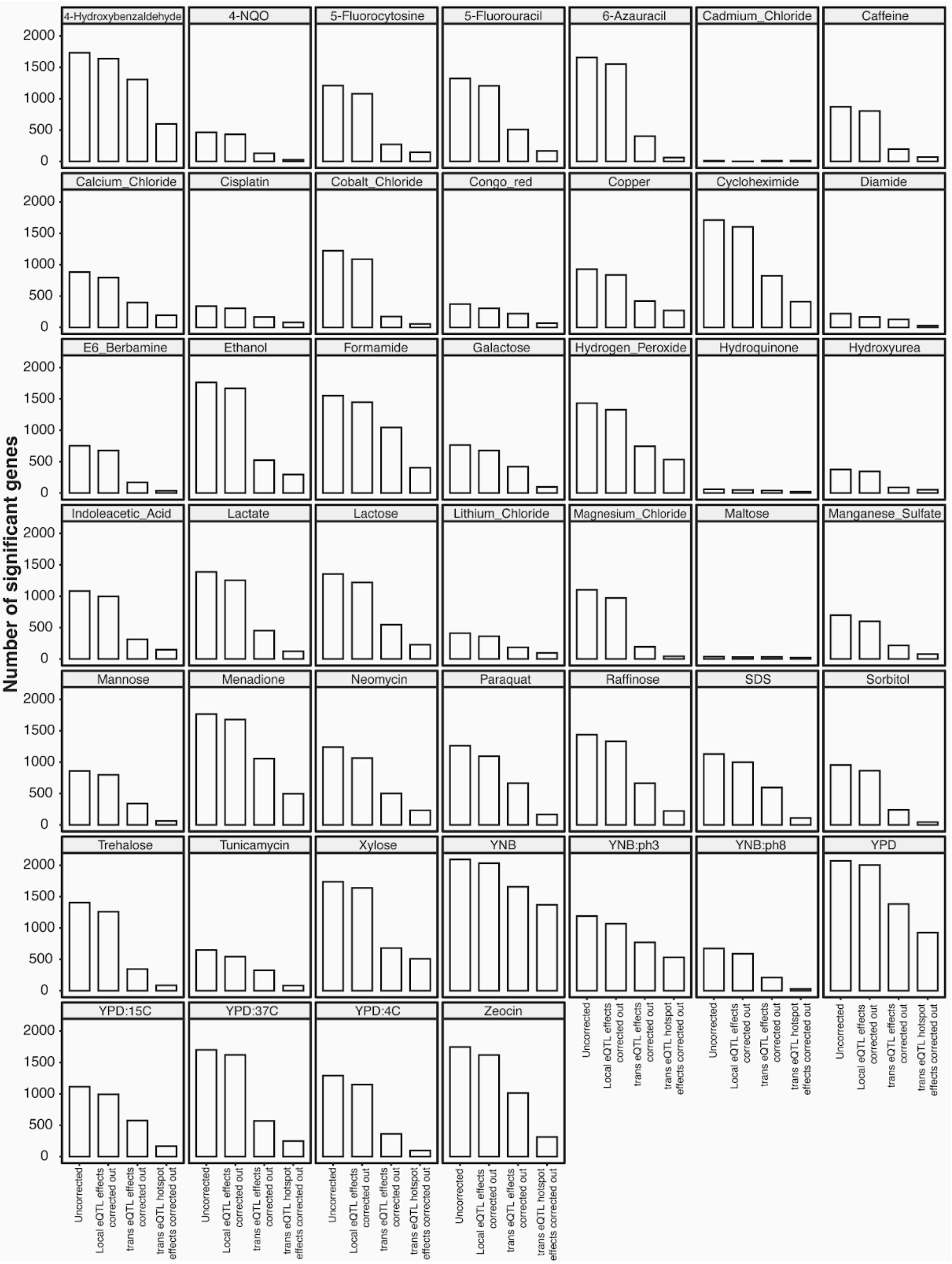


Supplementary Figure 11: Number of genes with significant genetic correlation before and after correcting for the effects of local eQTLs, *trans*-eQTLs and *trans*-eQTL hotspots. The 'uncorrected' category contains genes with significant genetic correlation at 5% FDR with at least one local eQTL and at least one *trans*-eQTL. Out of these genes, the number of genes with significant genetic correlation at nominal $p < 0.05$ after correcting out the effects of these genes' local eQTLs, their *trans*-eQTLs, and the 102 *trans*-eQTL hotspots is represented.

### Supplementary tables

Table S1: *Summary of growth traits* - Table containing the 46 growth conditions studied in Bloom et al., along with description of the base media used in these conditions, the number of segregants with finite growth measurement in Bloom et al., the total number of gQTLs mapped for each of these conditions by Bloom et al., the total variance explained by these gQTLs, and a description of the nature of the medium.

Table S2: *Heritability estimates and the number of genes with significant genetic, QTL effect, and hotspot effect correlations* - Table containing the number of genes with significant genetic, QTL effect and hotspot effect correlations for each of the 46 conditions at different significance thresholds, and the different heritability measures computed in this paper for these conditions.

Table S3: *Genetic, QTL effect, and hotspot effect correlation coefficients and colocalization information for 5643 genes and 46 growth conditions* - Table containing the genetic correlation, QTL effect and hotspot effect correlation coefficients and p-value for each of 5643 genes across the 46 growth conditions. The table also contains information about the eQTLs and gQTLs and their colocalization status for gene / trait pairs whose eQTLs and gQTLs were considered in our colocalization tests.

Table S4: *Number of overlapping and pleiotropic eQTLs for the 188 gQTLs considered in our colocalization tests*

Table S5: *Comparison of QTL effect and hotspot effect correlation results with the genetic correlation results* - Table containing the comparisons of the (i) correlation coefficients, and (ii) lists of significantly correlated genes, between the QTL effect correlations and genetic correlations analyses (Sheet 1) and from the hotspot effect correlations and genetic correlations analyses (Sheet 2) for each of the 46 conditions.

Table S6: *GO term enrichment results* - Table containing the results for enrichment of 163 GO-slim terms in the lists of genes with significant genetic correlations at 5% FDR (Sheet 1), significant QTL effect correlations at 20% FDR (Sheet 2) and significant hotspot effect correlations at 5% FDR (Sheet 3).

Table S7: *Mediation results* - Table containing the tests for mediation of the *IRA2* hotspot's effect on growth in hydrogen peroxide by expression of each of 1240 gene targets of the *IRA2* hotspot. For each gene, whether its expression is regulated by Msn2p is also indicated.

Table S8: *GO term enrichment results for significant mediators of IRA2 hotspot effect on growth in hydrogen peroxide* - Table containing the results for enrichment of 163

GO-slim terms in the 380 genes whose expression significantly mediates the *IRA2* hotspot’s effect on growth in hydrogen peroxide.
